# Identification of Specialized tRNA Expression in Early Human Brain Development

**DOI:** 10.1101/2025.01.22.633973

**Authors:** Alex L. Bagi, Todd M. Lowe, Sofie R. Salama

**Author notes:** **Lead Contact** Sofie R. Salama. **Corresponding author** Todd M. Lowe, Sofie R. Salama.

## Abstract

Transfer RNAs (tRNAs) are traditionally known for their role in protein translation, yet recent discoveries highlight their broader functions in gene regulation, particularly through tRNA-derived small RNAs (tDRs). Previous studies demonstrated the importance of one unique ArgUCU tRNA isodecoder in mouse neural development, yet potential function(s) of tDRs derived from this and all other tRNAs remain largely unexplored in human brain development. In this study, we employed cerebral cortex organoid models and AlkB-facilitated RNA methylation sequencing (ARM-seq) to profile tDRs across distinct stages of early human cerebral cortex development. Our analysis reveals dynamic expression patterns of diverse tDR groups derived from a wide range of isodecoders, with several distinct groups showing neural-specific expression. Computational analyses of these tDRs identified sequence motifs in over-represented tRNAs and enrichment for particular RNA modifications, giving initial clues to traits that define the pool of tDRs enriched during neural development. This expanded catalog of tDRs provides a framework for future studies on tRNA function in brain development and offers a deeper understanding of the complexity of tDR dynamics in neural differentiation.

## Introduction

A growing body of evidence suggests that tRNAs and their derived small RNAs (tDRs) play important regulatory roles that go beyond protein translation. Some of these extra-translational roles include global regulation of RNA silencing, immune system dysregulation, early developmental and cancer progression regulation.^1–5^ tRNA-derived small RNAs, in particular, have demonstrated roles in gene regulation and disease.^6–9^ Evidence suggests tDRs are also regulating cell homeostasis and differentiation via translation initiation/inhibition leading to altered cell-fate trajectories.^10^ Specific angiogenin-induced tDRs also have the ability to form into G-quadruplexes, which have neuroprotective roles in motor neurons, affecting the pathogenesis of diseases such as amyotrophic lateral sclerosis.^11,12^ A 5’ tDR from tRNA-Gly-GCC was recently shown to increase histone gene transcription in murine and human embryonic stem cells through its regulation of the U7 snRNA.^13^

Defects in tRNA modifications have also been associated with neurological diseases.^14^ For example, *FTSJ1*, which is a non-syndromic X-linked intellectual disability (NSXLID) gene, acts as a tRNA 2′-*O*-methyltransferase at two positions in the anticodon loop. Loss of *FTSJ1* function leads to decreased translation efficiency of UUU (Phe) codons, which are enriched in genes related to brain/nervous functions.^15^ Pseudouridylation of tyrosine tDRs via PUS7 has also been shown to selectively inhibit aberrant protein synthesis, and when altered leads to leukaemic transformation.^16^ Further, the bioavailability of specific amino acids such as arginine, glycine, and selenocysteine have been known to play roles in neurological function, disorders, and development, which may be influenced by insufficient levels of charged tRNAs or their associated tDRs.^17–19^ Recently, neurological disease-associated TRMT1 mutations were shown to cause a decrease in serine and tyrosine tRNAs due to loss of m2,2G26 modification, directly connecting aberrant tRNA modification with impaired neural development.^20^

Spatial- and time-dependent expression of protein-coding genes is an essential function for cell patterning and differentiation of neural tissues in mammalian neurodevelopment, but the roles of tRNAs and tDRs in this process have only recently been explored. A landmark study found that tRNAs produced by a single copy, deeply conserved arginine tRNA gene (tRNA Arg-TCT-4-1; *n-Tr20* in mouse) are abundant almost exclusively in the brain, and a single point mutation in the gene impairs the tRNA’s maturation, leading to severe post-natal neurodegeneration in the absence of ribosome release factors.^21^ Furthermore, loss of this specific tRNA leads to changes in seizure susceptibility and mTORC1 suppression, resulting in altered neurotransmission.^22^ Recent studies have found several instances where tDRs may also play roles in neurological diseases (reviewed in Qin et al. ^23^), but tDRs derived from Arg-TCT-4 have not been examined.

In this study, we sought to explore the expression patterns of tRNA fragments during early human brain development. We employed AlkB-facilitated RNA methylation sequencing (ARM-seq) ^24^ to enable a comprehensive analysis of this understudied class of small RNAs. ARM-seq uses the *Escherichia coli* dealkylating enzyme AlkB to remove a number of the most common tRNA modifications that often cause reverse transcription to terminate before reaching the end of a tRNA transcript. This and other methods, such as full-length tRNA specific QuantM-tRNA-seq ^25^, and YAMAT-seq ^26^, or tRNA and fragment gathering PANDORA-seq ^27^, MSR-seq ^28^, and hydro-seq ^29^, yield a much more complete picture of tRNAs and/or tDRs than standard small RNA sequencing methods. Furthermore, with a specialized tRNA analysis pipeline such as tRAX (tRNA Analysis of eXpression) ^30^, tDRs can be sorted into more complex categories than the general legacy naming groups ^31,32^ (e.g., 5’ and 3’ halves).

We applied ARM-seq to cerebral cortex brain organoids, an established model for early events in fetal brain development ^33–36^, to profile tDRs across various stages of development from pluripotent stem cells to the formation of excitatory projection neurons of the developing cortical plate.^37–39^ This approach enabled the identification of previously unknown brain-associated tDRs expressed during human cortical neurogenesis. We observed that, in addition to specific tDRs derived from the previously studied brain-specific Arg-UCU tRNA, a much wider array of tDRs exhibited changes, both in abundance and processing patterns. To further explore these dynamics, we employed dimensionality reduction, clustering, and classification techniques to group the tDRs we identified based on fragment properties. Among the multiple sub-classes of neural tDRs found, those derived from selenocysteine, alanine, and glycine tRNAs were most notable. Collectively, they show conservation bias at a number of positions coincident with specific tRNA modifications, including m^2^ G26, previously shown to be critical for normal brain development.

## Results

### Validation of organoid model and RNA profiling

Human induced pluripotent stem cells (GM12878-c305 iPSCs) were grown into cerebral cortical organoids as a model to study tDR changes in early human brain development. A standard time course of culturing conditions was used to achieve neural induction and organoid growth **(Figure 1A)**. Cells were aggregated into embryoid bodies and differentiated into cortical tissues over a period of 10 weeks (see Methods) to span the emergence of key cell types in the course of dorsal cortex development^37^ and sampled at four time points: day 0 (D0 - stem cells); day 14 (D14 - neural epithelium and radial glia neural progenitor cells), day 35 (D35 - emergence of deep layer neurons) and day 70 (D70 - continued generation of cortical projection neurons, emergence of outer radial glia). D70 organoids expressed expected neural cell-type markers using immunofluorescence staining (IF-staining) when compared against D0 stem cells **(Figure 1B-D)**. Circular structures known as neural rosettes (early neural tube-like formations) were observed throughout the perimeter of the organoids when stained with antibodies for PAX6 (a radial glia neural stem cell marker), and vimentin (a radial glial fiber marker), confirming the presence of radial glia neural stem cells **(Figure 1B)**. Furthermore, these PAX6+ neural rosettes were surrounded by cells staining for TBR2 (an intermediate neural progenitor marker), and CTIP2 (a deep layer neuron marker) **(Figure 1D)**. The presence of different neural subtypes increased over time as the organoids developed and increased in size, validating expected dorsal forebrain organoid morphology and developmental trajectory.

**Figure 1:**
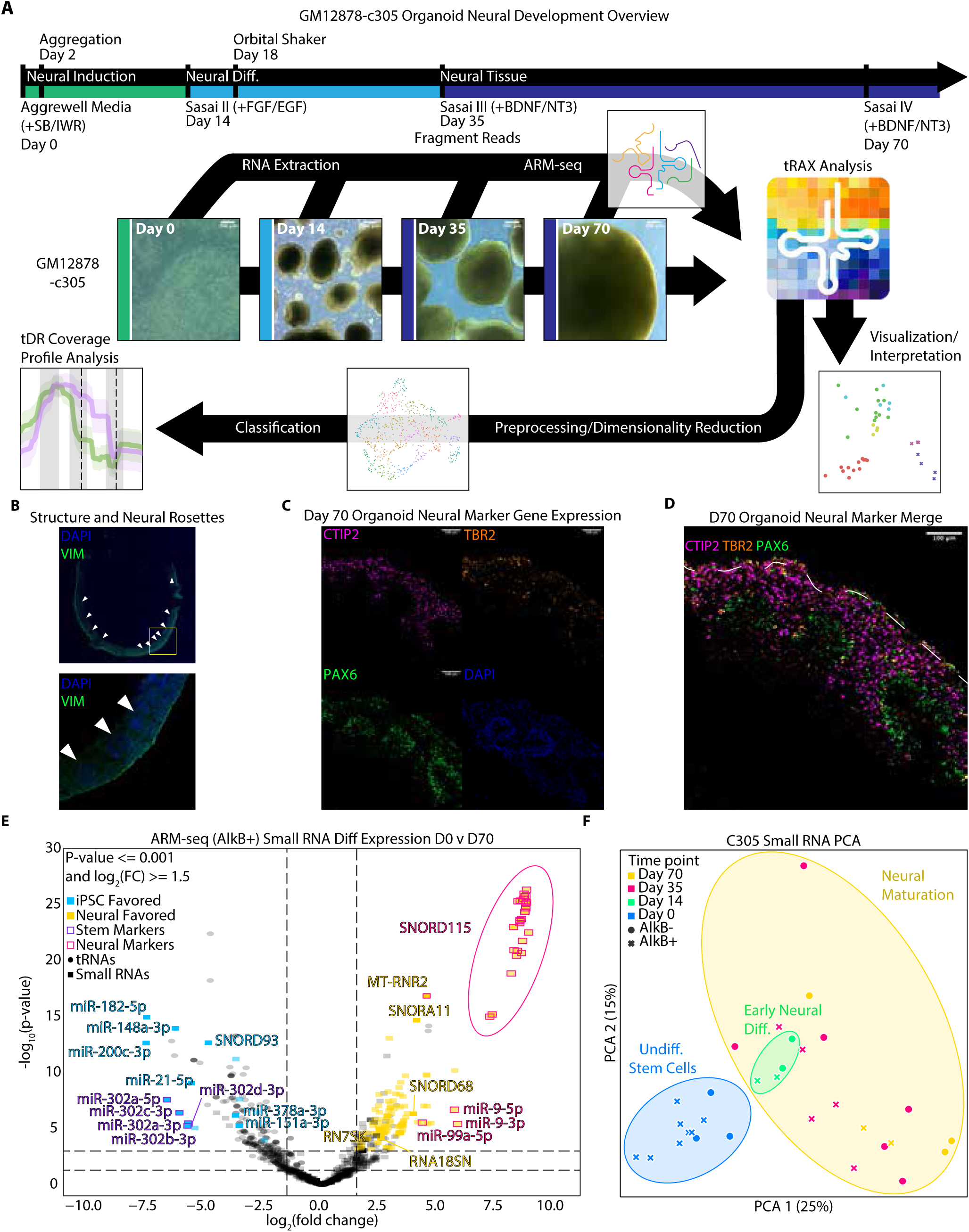
Validation of organoid model and RNA profiling. **(A)** Graphical abstract of the study depicting media and small molecule drug changes across the organoid developmental time course. Representations of organoids and relative size at crucial time points are shown with 500 μm scale bars. RNA collection, tRNA sequencing, analysis pipeline, and classification are also depicted. **(B)** D70 organoid stains using vimentin (VIM) and DAPI are used, showing cytoarchitecture of the organoid. Emphasis placed with white arrows showing locations or neural rosettes (early endotube-like formation) found throughout the perimeter of the organoid stains. **(C)** Antibody stains PAX6, a radial glial marker, CTIP2, a deep layer neuron marker, and TBR2, an intermediate neural progenitor marker, show neural marker gene expression with PAX6 staining the inside of neural rosettes, TBR2 surrounding the rosettes and CTIP2 encompassing the exterior of the organoid. **(D)** Neural marker stains were merged with a dotted white line for the exterior perimeter of the organoid. **(E)** Volcano plot showing D0 vs. D70 log_2_ fold-changes in the abundance of small RNAs against −log_10_(p-value) with tRNAs shown for scale. Targets of particular interest are labeled with horizontal dotted lines at p-value=0.05 and p-value=0.001 and vertical dotted lines at 1.5 fold difference. Small RNAs highly expressed in either stem or neural cells across multiple conditions (D0 vs. D14, D0 vs. D35, or D0 vs. D70) were highlighted in blue or yellow, respectively. Small RNAs known as stem or neural markers were highlighted in purple or magenta as well. **(F)** PCA plot of all ARM-seq samples for all measured small RNAs across organoid time points.

To observe high-resolution tDR dynamics in this neural development model, we performed ARM-seq small RNA sequencing ^24^, followed by differential expression analysis using tRNA Analysis of eXpression (tRAX) ^30^ for samples representing the differentiation time course (D0, D14, D35, D70). We validated that our small RNA sequencing reflected neural differentiation with multiple classes of small RNA markers known to have specific expression in neurons or stem cells **(Figure 1E)**. Stem-associated mir302 family miRNAs were more highly expressed in D0 stem cells, whereas neural-associated miR-9 and miR-LET7 (miR-99a) family miRNAs were more highly expressed in the D70 organoids **(Table S1)**.^40–43^ MT-RNR2, a mitochondrial ribosomal RNA found to play a neuroprotective role, was also upregulated in the organoids relative to stem cells.^44–46^ Many SNORD115 small nucleolar RNAs, known to be induced by neuronal differentiation^47^, were also highly expressed in the later time points, confirming expected small RNA transcriptional changes during the organoid time course experiment. Principal Components Analysis (PCA) of small RNA read counts across all time points **(Figure 1F)** confirmed that biological replicates and adjacent time points cluster closest to one another.

### Identification of complex and dynamic tDR profiles during organoid differentiation

The relative expression of the major classes of small RNAs was assessed across the developmental time course, and tRNA reads increased in abundance with AlkB treatment, as expected for the most highly modified class of small RNA **(Figure 2A; Figure S1)**. At a very high level, it is clear that a complex mixture of tDRs derived from both 5’ and 3’ tRNA ends changed, with Ala, Asn, iMet, Phe, SeC, and Ser showing increases of more than two-fold between D0 and D70 **(Figure 2B),** with the greatest relative expression change between D0 and D35. In contrast, more than 40% of Glu, His, Lys, Thr, and Tyr tDRs decreased over the full time course, with His and Tyr showing the most immediate decreases between D0 and D35 conditions.

**Figure 2:**
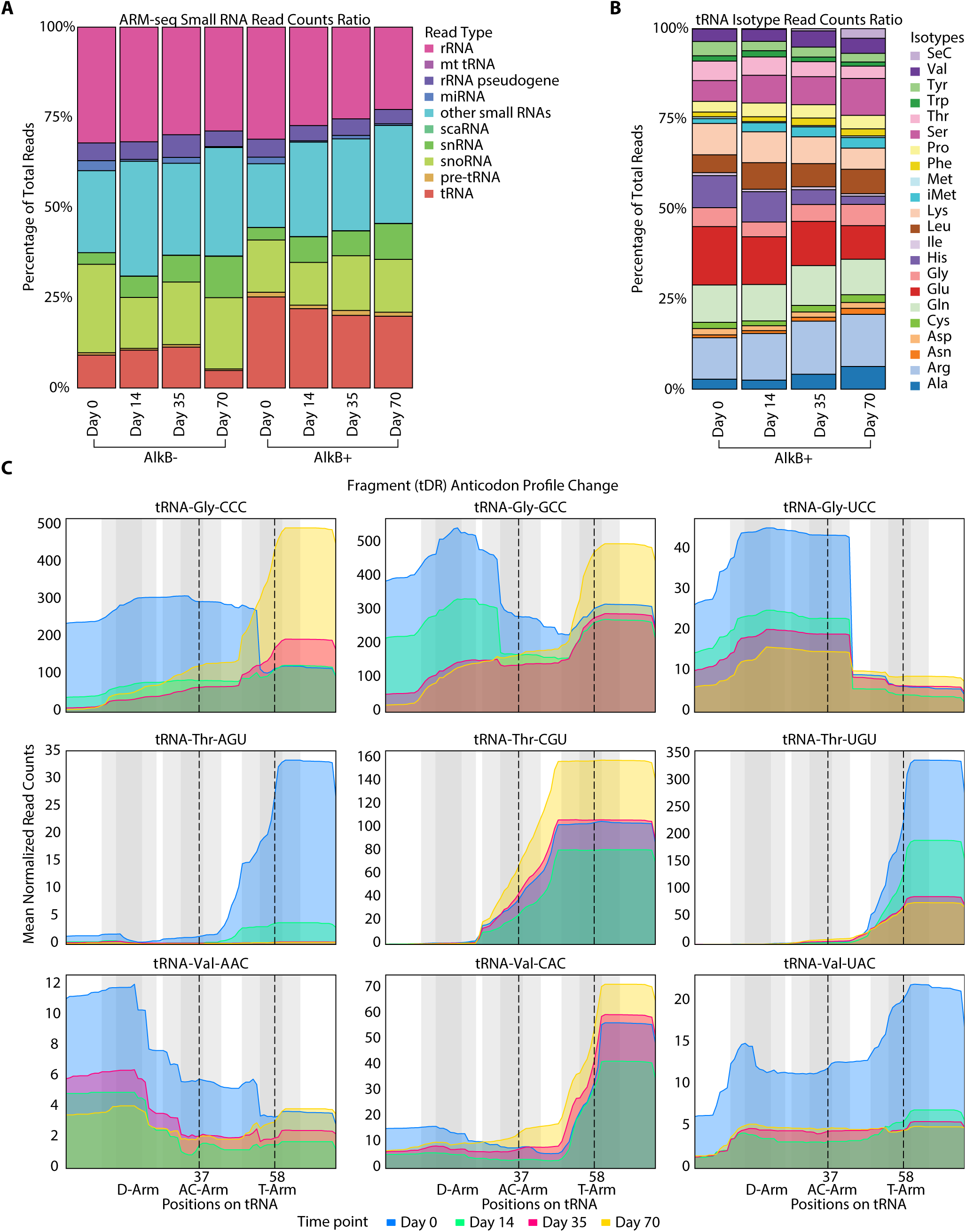
Identification of complex and dynamic tDR profiles during organoid differentiation. **(A)** Human small RNA read count distributions showing relative ratio of small RNAs across organoid time course, sequencing type, and AlkB treatment. **(B)** Human tDR isotype read counts (total and fragment) distributions showing relative ratio of tRNA isotypes across organoid time course, sequencing type, and AlkB treatment. **(C)** Coverage profiles for the Gly, Thr, and Val isotype groups showing max expression across AlkB+ samples per position within each isotype across organoid time points. D-loop, anticodon-loop, and T-loop shown in gray, with lines added at specific coverage breakpoints in each plot.

However, strictly measuring total fragment counts obscures a much more biologically complex picture of the molecular changes in tDR processing. Coverage profiles for the most dynamic isotypes (Gly, Thr, and Val) tDRs show remarkable changes in expression and diversity in tDRs between isoacceptors across developmental time points, with 5’ versus 3’ fragments, of different lengths, showing very different trends for tDRs derived from tRNAs decoding the same amino acid **(Figure 2C, Sup Figure S2)**. For example, different Gly 5’ fragments evident in D0 samples (Figure 2C, Gly-CCC, Gly-GCC, Gly-UCC, blue coverage) diminished over later time points, although at different rates among the three isoacceptors. Notably, Gly-GCC 5’ tDR has been shown to control histone expression in pluripotent stem cells.^48^, consistent with its prominence in our data. In contrast, 3’ tDRs derived from Gly-CCC and Gly-GCC increase in abundance as the cells differentiate. For Thr-derived tDRs (Figure 2C, three middle plots), shorter 3’ fragments (ending near position 58 in the T-loop) decrease for Thr-AGU and Thr-UGU isoacceptors, but longer Thr-CGU-derived tDRs (ending in the variable stem) *increase* expression over time, suggesting that the differences in sequence and/or modifications, particularly at the tDR ends, could play a role in the specificity of processing and stability of these fragments. Val tDRs (Figure 2C, three bottom plots) have the widest variety of tDR coverage profiles with a distinct pattern for each time point and isoacceptor. In addition to these, many other isotypes have striking changes across the time course, showing complex dynamics in abundance and fragment types **(Figure S2A).**

### Identifying neural-specific versus stem-like tDRs

Individual tDRs were examined to identify and rank the most “neural-specific” fragments emerging during the neural differentiation time-course**(Table S2)**. The expression of D0 vs. D70 tDRs was compared for all tDRs and pre-tRNA type reads, showing very different distributions for iPSC-favored versus neural-favored tDRs **(Figure 3A)**. Between D0 and D70 samples, 27 tDRs were found to have a significant (p-value <= 0.05) and log_2_(fold-change) greater than 1.5 increase in expression in the cerebral cortex organoids relative to undifferentiated stem cells **(Table S2)**. The top 15 neural- and top 15 stem-expressed tDRs between D0 and D70 were visualized by log fold change **(Figure 3B),** p-value, and normalized read counts **(Figures S3A,B)**. Among the top rank-expressed neural tDRs, several isotype families appeared multiple times: Ala (4), Arg (3), Gly (2), and Leu (3).

**Figure 3:**
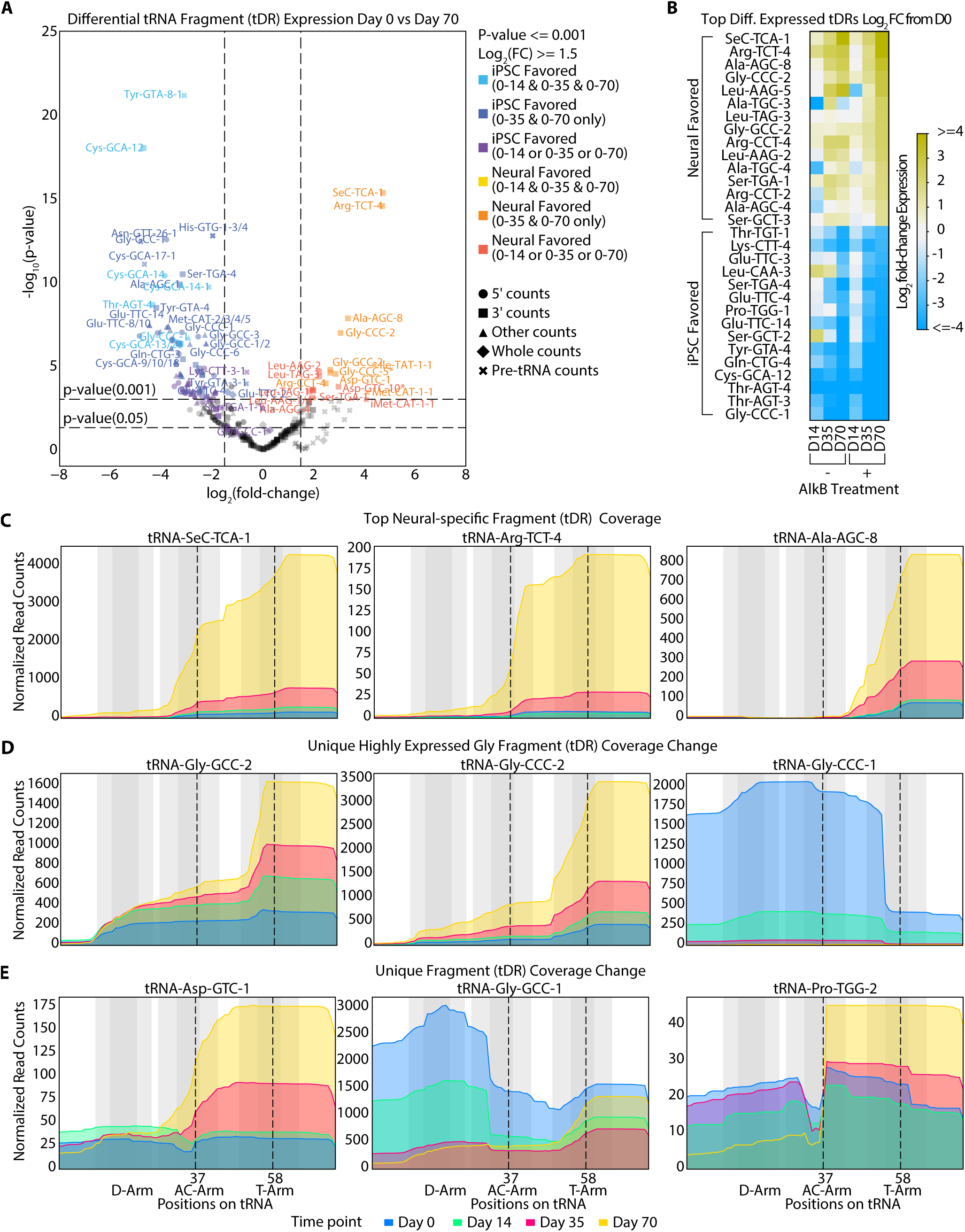
Identifying neural-specific versus stem-like tDRs. **(A)** Volcano plot showing D0 vs. D70 log_2_ fold-changes in the abundance of tDRs against −log_10_(p-value) with 3’, 5’ and internal fragment counts and full-length and pre-tRNA counts. Targets of particular interest are labeled with horizontal dotted lines at p-value=0.05 and p-value=0.001 and vertical dotted lines at 1.5 fold difference. tDRs highly expressed in either stem or neural cells across multiple conditions (D0 vs. D14, D0 vs. D35, or D0 vs. D70) were highlighted in shades of blue or yellow, depending on how many conditions they appeared in. All samples were batch-corrected and normalized using DESeq2 ^61^ to account for differences in sample read counts. **(B)** Heatmap showing the top 15 neural and top 15 stem-favored tDRs across all conditions. Names in orange have less than 80 uniquely mapping read counts. **(C-E)** Coverage profiles for the top 3 highly expressed tDRs over organoid neurogenesis **(C)**, highly expressed stem and neural Gly tDRs **(D)**, and unique tDRs with coverage changes **(E)**. D-loop, anticodon-loop, and T-loop shown in gray, with lines added at specific coverage breakpoints in each plot.

In our data, nearly all tDRs that increase after neural differentiation are 3’ end-counts (3’ fragments that terminate at, or within 10 bp of the CCA tail), with several pre-tRNA counts also being detected. The top 15 neural- and top 15 stem-expressed tDRs between D0 and D70 were compared **(Figure 3B)** to determine overall significance and expression **(Figure S3A,B)**. While 5’ end-counts (5’ fragments that terminate within 10 bp of the tRNA start site) are less common when compared to 3’ end-counts, they tend to appear in D0 samples and diminish entirely after neural differentiation in both D35 and D70 time points. With many 3’ tDRs being expressed at D35 and D70, there is also great diversity in the shape of fragment coverage with drop-offs of coverage at different conserved tRNA positions. This suggests differential modifications of the mature tRNA from which the tDR is derived since mature tRNA modifications often act as signals for cleavage creating tDRs.^49^

The diversity of the top three most neural-expressed tDR 3’ fragments was explored in detail **(Figure 3C)**. While coverage drop-off (5’ ends of tDRs) commonly occurred near position 58, other 5’ start positionsdiffered between tRNA isodecoders. tRNA-SeC-TCA-1 and tRNA-Arg-TCT-4 derived tDRs, for example, are longer, with 5’ ends occurring within the anticodon loop, making these tDRs match the footprint of traditional 3’-tRNA halves. Additionally, while both tRNA-Arg-TCT-4 and tRNA-Arg-TCT-1 tDRs increase in abundance over time, the former has a severe drop-off in coverage around position 40, and the latter persists into the D-arm before dropping off **(Figure S3C)**. tRNA-Ala-AGC-8-derived tDRs, however, had distinct 5’ ends between the anticodon and T-loop **(Figure 3C)**. tRNA-Gly-GCC-2- and tRNA-Gly-CCC-2-derived tDRs also had distinct fragment end-points appearing in the top 10 most neural-expressed tDRs, in surprising contrast to tRNA-Gly-CCC-1, one the most stem-expressed tDRs. **(Figure 3D)**. Interestingly, both neural-specific Gly tDRs (tRNA-GCC-2/CCC-2) were 3’-specific whereas tRNA-Gly-CCC-1 is highly 5’ biased with 3’ ends occurring before position 58. tRNA-Sec-TCA-1, tRNA-Arg-TCT-4, and tRNA-Ala-AGC-8 were validated as increasing in full-length tRNA expression across neural differentiation via Northern blots **(Figure S3D)**. From these data, we observe that when examined individually, most tDRs maintain roughly the same fragment type and endpoints over the time course, but change in abundance, suggesting source tRNA abundance or tDR stability changes but not necessarily the tDR processing pathway.

A few tRNAs showed pronounced changes in the tDR fragments generated over the developmental time course. tRNA-Asp-GTC-1-derived tDRs look similar in shape and expression in the first two time points but gain a large increase in 3’ fragment expression in the last two **(Figure 3E)**. tRNA-Gly-GCC-1-derived tDRs have a contrasting pattern with the diminishment of the 5’ end tDR coverage as the organoids reach the last two time points. tRNA-Pro-TGG-2 tDRs exhibit a mix of both patterns, with a decrease in 5’ fragments paired with an increase in 3’ fragments over time. Since excitatory neuron formation increases between D35 and D70, 3’ fragments may be associated with these neurons. tRNA-Arg-TCT-2 and tRNA-Ser-GCT-5 tDRs form a unique group that shows higher expression in D0 and D70 relative to D14 and D35, which may suggest lower expression in neural progenitor cells **(Figure S3E)**.

### Multidimensional analysis reveals tDR clusters associated with neural vs. stem states

To assess the full diversity of unique tRNA expression, clustering and classification of fragment profiles split across different time points was performed. This was done by aggregating read coverage, read-starts/ends, deletions, and misincorporation information (suggesting modified positions) and clustering the combined dataset using UMAP (Uniform Manifold Approximation and Projection) followed by classification via HDBSCAN.^50,51^ The resulting clusters were then manually annotated into neural and stem categories as described in the methods. This revealed a highly complex, diverse set of tRNAs clusters, each comprising multiple isotypes, and neural-stem expression levels with no bias in time point observed **(Figure 4A)**. Three major groups of HDBScan clusters were identified (consisting of 21 specific subgroups) labeled A (stem), B (neural), and C (neutral) as well as the HDBSCAN unannotated category (U) for tRNAs that were too hard to class into a single cluster. Spatially clusters tended to correlate with isotype and neural/stem favored tRNAs with no observable bias in time point. We next explored the relationship between isotype and cluster to see if genomic traits could also be correlated to tDR expression.

**Figure 4:**
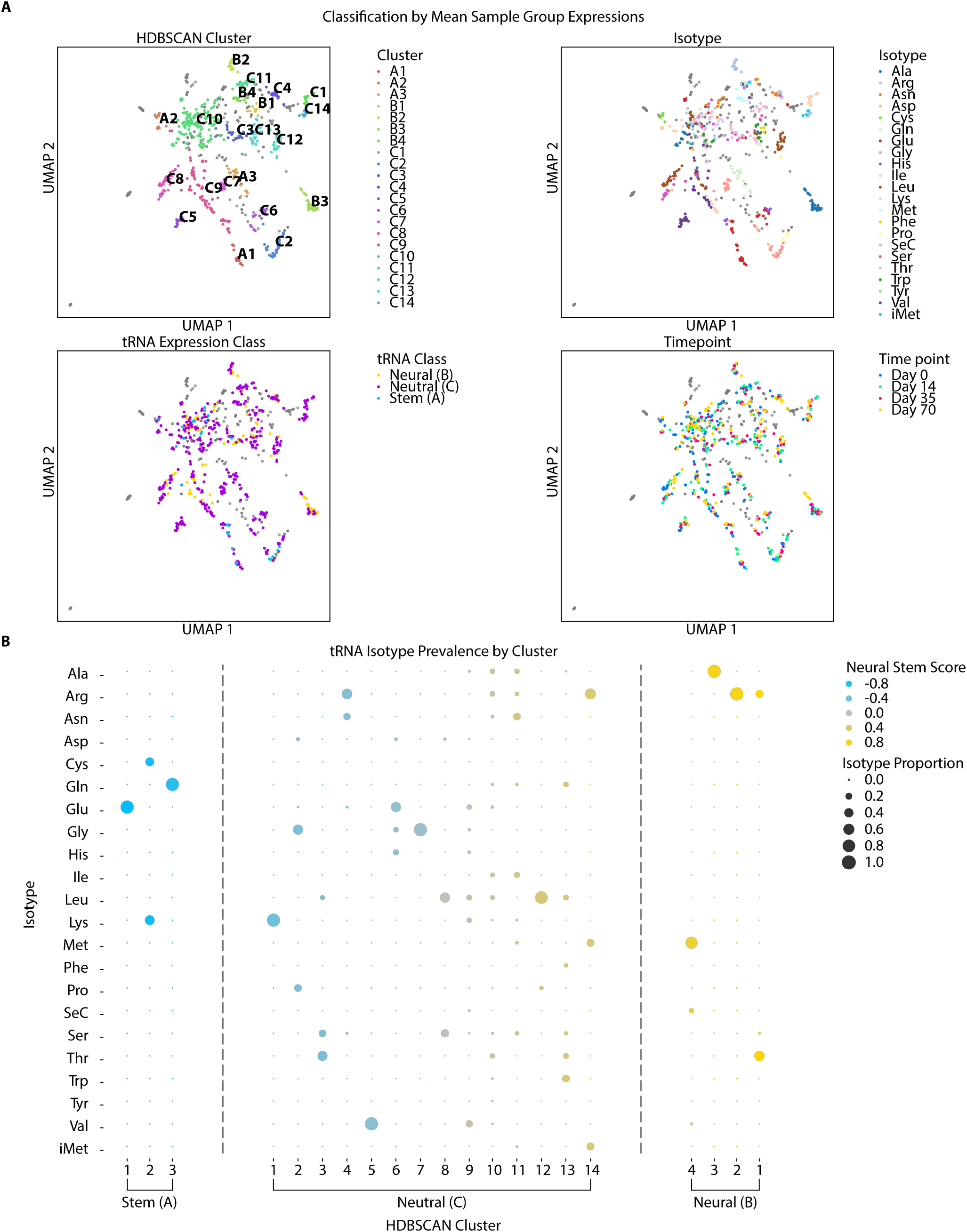
Multidimensional analysis reveals tDR clusters associated with neural vs. stem states. **(A)** UMAP projections of tDR sequencing profiles, using unique coverage of tDR 3’ and 5’ read ends, read alignment deletions (see methods), and mismatched bases, by position in the tRNA, using HDBSCAN clustering and manual annotation of stem (A*n*), neural (B*n*), and neutral (C*n*) classes. tDR fragment profiles that could not be confidently annotated into clusters by HDBSCAN were masked (grey) and not included in further analysis. **(B)** Dot plot representation of the HDBSCAN clusters with dot size correlating to the proportion of isotype found in each cluster and the color based on the mean neural stem score (as defined in methods) with vertical lines at the top and bottom 25th percentiles defining the groups.

The clusters were often dominated by specific tRNA isotypes with several clusters containing greater than 50% of specific isotypes: A1 (Glu), A2 (Gly), A3 (Gln), C1 (Lys), C2 (Asp), C3 (Leu), C5 (Val), C6 (His), C7 (Gly), C8 (Ser), C12 (Leu), C14 (Arg), B1 (Ala), B2 (Arg), and B3 (Ala) **(Figure 4B)**. The clusters correlate with distinct isotypes **(Figure S4A)**, AlkB treatment **(Figure S4B)**, and neural expression patterns **(Figure S4D)** suggesting that specific RNA sequences drive these clusters even though only positional expression information and not base identity was used in the clustering. In contrast, Cluster C10 and C11 both have diverse isotype compositions and seem to correlate more closely with the unannotated (U) category. This can likely be explained by low mean read coverage in clusters C10 and C11 relative to other clusters making them hard to annotate **(Table S3)**. AlkB conditions were generally diverse with a slight preference for AlkB+ conditions in clusters with higher neural expression. We checked whether subtle genomic features of tRNAs such as tRNAscanSE covariance, HMM, and secondary structure scores (Infernal - Inference of RNA Alignments) correlated with cluster assignments and found that scores tended to match within clusters **(Figure S4C)**. These scores measure how much each individual tRNA sequence matches its canonical isotype and is consistent with our observation that individual isotypes define certain clusters. This analysis suggests that differential expression and tDR formation are influenced by these genomic features but most likely they are not the only driving factor.^52,53^

To further examine the relationship between isotype and cluster, we looked at Arg specific tDRs. Disregarding U, C10, and C11 (due to aforementioned indiscernibility) Arg tDRs are split across B1, B2, C4, and C14 clusters all residing in the same spatial area as one another **(Figure 4A; Table S3)**. Interestingly, Arg-TCT-4 shows up in both B1 (neural) and C10 (neutral) clusters with D70 Arg-TCT-4 appearing in B1. This is most likely due to the extreme change in expression and coverage after neural differentiation with lower read coverage harder to discern **(Figure 4A; S4D)**. This indicates that other “neutral” tDRs appearing in C clusters may also be neurally associated but can be hard to discern with low read coverage. B1 clusters also contain Arg-TCG-1 and Arg-TCG-5 tDRs, with B2 clusters comprised almost entirely of Arg-CCT tDRs, all tRNAs that increase moderately over differentiation. Interestingly Arg-TCT-1 (which also increases across neural differentiation) appears in C10 (neutral) and U (unannotated) clusters with D70 appearing in C10, this mimics the change over the time course seen in Arg-TCT-4 albeit from U to C10 rather than C10 to B1. While both Arg-TCT-1 and Arg-TCT-4 increase in neural expression, the former has less of an expression increase than the later, and they express entirely different fragments. This revealed that Arg tDR expression as a function of neural expression is correlated to specific tRNA transcripts rather than specific isotypes and subtle changes to genomic sequence correlate with broad effects in the type and expression levels of tDRs.

### Sequence motifs and modifications associated with neural tDR clusters

To further understand what causes specific neural tDR shape and expression we looked at tRNA modifications and their enzymes and how they correlated with underlying characteristics of the parent tRNA. Modifications were derived for all human tRNA transcripts from data in Zhang et al. ^20^ (see methods), and were projected onto the clusters showing that specific modifications such as acp^3^U20, m^3^C20, and m^1^I37 correlate strongly with just neural clusters (B), and acp^3^U20a, m^2^ G26, m^3^C32, and I34 correlate with neural and neutral (B and C) clusters **(Figure 5A)**. Inosine (at position 37) has been associated with intellectual disabilities, and microcephaly and is added by ADAT3 in humans. m^2^ G (a modification found at position 26 in Arg-TCT-4) has been associated with intellectual disabilities and microcephaly via TRMT1.^54,55^ m^1^A58 is a ubiquitous modification found in nearly all tRNAs and is consistently expressed across all groups. While not present in our modification data, PUS3 and NSUN2 are known to play a roles in pseudouridylation and m^5^C addition at positions 39 and 40, respectively, and are both tied to intellectual development.^56–58^ We found that PUS3 and NSUN2 increased while ADAT3 decreased over a five week time course of human embryonic stem cell derived cortical organoids using data from Fiddes et al. ^38^ **(Figure S4E)**.

**Figure 5:**
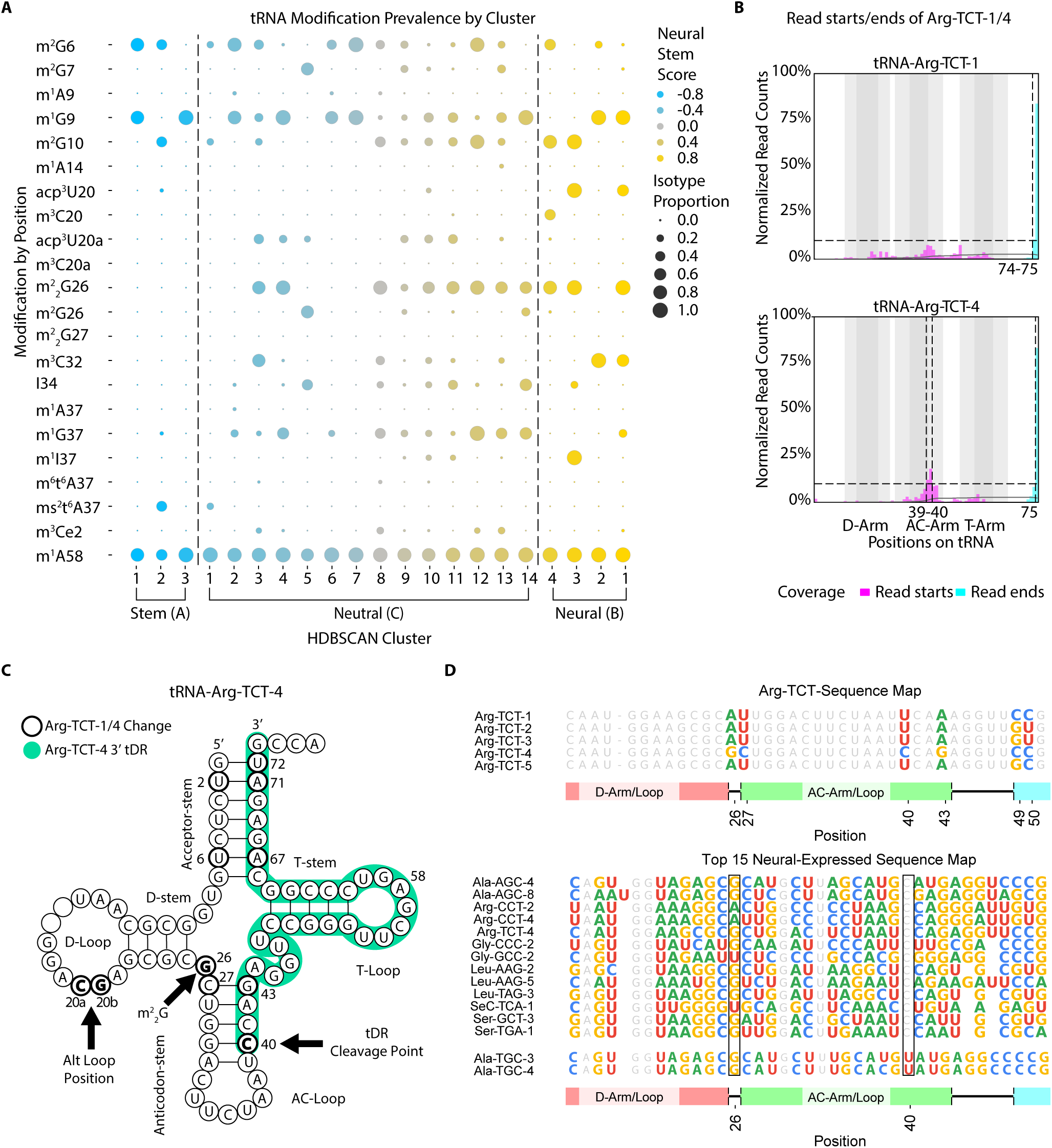
Sequence motifs and modifications associated with neural tDR clusters. **(A)** Dot plot representation of the HDBSCAN clusters with dot size correlating to the proportion of modification at positions found in each cluster and the color based on the mean neural stem score (as defined in methods) with vertical lines at the top and bottom 25th percentiles defining the groups. **(B)** The percentage of 3’ and 5’ read ends of Arg-TCT-1 and Arg-TCT-4 tDRs. Positions with greater than 10% of relative normalized fragment read-starts/ends indicated with vertical dotted lines. **(C)** Secondary structure of Arg-TCT-4 showing divergence with Arg-TCT-1, key differences in bold and highlighted with arrows. **(D)** Sequence map of Arg-TCT tRNAs and top 15 most neural-expressed tDRs.

Sequence alignments of the tRNAs in the neural clusters (B) revealed conservation of tRNA sequence at positions 26, 30, 32, and 40 **(Figure S5A)**. This includes bases in close proximity to the cleavage at position 40. We explored whether sequence motifs (relative to background frequency) played a role in clusters as they have potential to affect tertiary structure formation. Motifs such as GGGG (1-4, 68-71), and CCCC (2-5, 60-63) and GCC were observed to be more common in neural (B) clusters **(Figure S5B)**. The motif GCAU (26-29) also indicates a higher frequency of G at position 26 which most likely corresponds to the higher frequency of m^2^ G26 found in the neural (B) clusters. AUG (28-30), GUA (39-41) and GCC (39-41 in neural, 40-42 in stem) indicated that the G-C pair between 30 and 40 occurs more frequently in neural groups but a G is more likely at position 40 in stem. With neutral (C) and unannotated (U) clusters comprising the majority of tRNAs found in the clustering, relative motif frequency was much closer to expected background.

Finally we also looked at the internal arrangement of the tDRs for both Arg-TCT-1 and Arg-TCT-4 by looking at the ratio of 3’ read ends (in relation to the tRNA structure) to 5’ read ends **(Figure 5B)**. In Arg-TCT-4 read starts are grouped around position 39 and 40 indicating an association with having a C rather than U at that position as found in other Arg-TCT isodecoders. Arg-TCT has five isodecoders, with Arg-TCT-4 having multiple unqiue bases relative to the others including a 20b position G as well as 2U, 20aC, 26G, 27C, 40C, 43G, 71A, and 72U **(Figure S5C)**. Further when we compared Arg-TCT-4 against the 15 top neural expressed tDRs we found conservation of C at position in 40 in all but Ala-TGC-3 and Ala-TGC-4 **(Figure 5D; Figure S5C)**. Interestingly Ala-TGC-3/4 both express more in D0 relative to D14 before increasing in D35 and D70 suggesting that they are active at different neural time points and might be part of a different neural expression pathway. Together these results define a shared set of features associated with tRNA genes giving rise to neural-specific tDRs.

## Discussion

tRNA gene expression has been previously examined in multiple neurological contexts ^22,59,60^, including neural cell lines and mature brain tissue, but none designed to capture key developmental transitions from human pluripotent stem cells to the early cerebral cortex and these studies did not focus on tDRs. This study unveils striking changes in tDR abundance and processing patterns during early human corticogenesis, the first requisite step towards identifying neural-specific tRNA processing patterns which may have key roles in brain development. We have identified many unique tDRs that change their expression profile and have categorized them. Furthermore, we developed a method that categorized these complex tDR patterns by clustering transcripts based on processing patterns, sequence, and modifications. This allowed us to look for commonalities in processing that might correlate with downstream function.

tRNA-Arg-TCT-4 has been previously identified in mouse as a neural specific tRNA with defects leading to truncal ataxia and severe neurodegeneration when position 50 has a C to T mutation.^21^ Here we show that Arg-TCT-4 derived 3’ tDR abundance increases across early human corticogenesis as well. This profile of brain organoids has also allowed for the discovery of new tDRs that are induced during neural differentiation, such as those from SeC-TCA-1 and Ala-AGC-8. Further, while most studies focus on single tRNA transcripts, using ARM-seq combined with cerebral brain organoids has allowed for the discovery of many tDRs with dynamic expression in early human embryonic development. These fragments each have unique breakpoints in their read coverage as well, indicative of a unique modification profile on a tRNA by tRNA basis. Intriguingly, the top hits for neural specific tDRs seem to share a coverage profile similar to one another, suggesting they may be generated by a common mechanism.

The profiles of these tDRs tended to fall into similar patterns that could be visually grouped together. For example, some of the top neural favored tDRs (Arg-TCT-4 and Ala-AGC-8) each have a 3’ fragmentation pattern that is visually unique, reflecting a combination of associated sequence factors that allows for the fragments to be classified into distinct clusters that are proximal to one another (B1 and C13). This suggests that advanced sequence characteristics beyond unique read coverage drive the grouping of neural expressed tDRs. The unique fragmentation patterns of highly-expressed neural tDRs, such as those derived from Arg-TCT-4 and Ala-AGC-8, are not simply a consequence of increased tRNA abundance. Rather, they represent a qualitatively different mode of tRNA processing, dictated by a complex interplay of sequence motifs, likely including specific modifications and the enzymes that catalyze them. These results strongly suggest that particular tDR isoforms, and the processes that generate them, are actively selected for during neurodevelopment. They also suggest that perturbing this modification program either through modification of the tRNA gene or the relevant modifying enzymes could be used to probe the functional relevance of these neural-specific tDRs.

While this approach was used in the context of human cerebral cortex differentiation, identifying and classifying tDRs can be used in a broader context. Understanding the relationship of tDRs in a more expansive way than 3’ and 5’ halves can lead to a better understanding of tDR formation overall and their roles in RNA processing beyond translation. Our study revealed a coordinated, dynamic remodeling of the tDR landscape during human corticogenesis. This is not merely a shift in abundance, but a precise orchestration of tRNA processing, generating specific tDR isoforms with distinct cleavage patterns. The clustering of these tDRs, driven by a combination of sequence features, modifications, and expression profiles, points towards functionally distinct classes. This work establishes a framework for dissecting the molecular mechanisms by which tDRs influence tissue development, opening new avenues for understanding the etiology of developmental disorders and diseases, suggesting potential therapeutic targets based on tRNA processing.

## Resource availability

### Lead Contact

Requests for further information and resources should be directed to and will be fulfilled by the lead contact, Sofie R. Salama.

### Materials Availability

This study did not generate new unique reagents.

### Data and Code Availability

All raw and processed sequencing data generated in this study have been submitted to the NCBI Gene Expression Omnibus (GEO; https://www.ncbi.nlm.nih.gov/geo/) under accession number GSE259250 (https://www.ncbi.nlm.nih.gov/geo/query/acc.cgi?acc=GSE259250). All computational methods are provided in a series of Jupyter notebooks available on GitHub (https://github.com/alba1735/organoidtRNAs/).

## Acknowledgments

We received technical support with microscopy from Benjamin Abrams, UCSC Life Sciences Microscopy Center (RRID: SCR_02113). For Cell culture we received support from the CIRM Major Facility Award to UCSC (FA1-00617) and the IBSC Stem Cell Culture Facility (RRID:SCR_021353). Additionally, Dylan Baer assisted with tissue culture and cerebral cortical organoid growth, and Liam Tran helped with IF-staining methods and troubleshooting. We also received support from Jonathan Howard and Kristof Tyigi for advice on experimental methods, technical assistance, and support in RNA library preparation and general laboratory methods. Finally Aidan Manning collaborated in the creation of the modification data file for analysis.

## Author contributions

A.B., T.L. and S.S. designed the experiments and interpreted the results. A.B. performed all experiments, conducted the data analysis, generated the figures, tables, code, and wrote the manuscript. T.L. and S.S. provided critical review and editing of the manuscript.

## STAR Methods

[Key Resources Table](https://docs.google.com/spreadsheets/d/11OfFiDkQ2-ptV1PpK6uTOA_eKLDWDJZrdKx1FjXR02I/edit?usp=sharing)

### Organoid methods

Human GM12878-c305 induced pluripotent stem cells (iPSCs) were generated from the GM12878 lymphoblastoid cell line (Coriell) using a proprietary episomal vector-based reprogramming method by Cellular Dynamics International (fujifilmcdi.com) and were confirmed to have a normal karyotype using the KaryoStat Karyotyping service (ThermoFisher). Undifferentiated cells were grown in feeder-free conditions on matrigel (Corning) with mTeSR Plus (Stemcell Technologies) or on vitronectin with Stem Flex media (ThermoFisher). Cerebral cortex organoids were generated using a protocol adapted from Kadoshima et al. ^36^. 10,000 cells per embryoid body were aggregated using AggreWell-800 plates in AggreWell media (Stemcell Technologies) supplemented with 10 uM Y-27632 rock inhibitor (Stemcell Technologies), and transferred to low attachment 6-well dishes (Corning) on day 2. These methods supplemented the respective media with 10 uM SB431542 (SB, Millipore), and 1 uM IWR-1 (Millipore) for the first 14 days of differentiation. The media was then changed to Sasai II media (DMEM/F12 + glutamax supplemented with N2) on day 14. At this point, organoids were supplemented with 10ng/mL beta fibroblast growth factor (bFGF) and 10ng/mL epidermal growth factor (EGF) to improve survival in Sasai II media. On day 18 the organoids were transferred to an in-incubator orbital shaker (100 rpm) where they remained for the duration of the experiment. After day 35, all cultures were grown in Sasai III media (Sasai II media supplemented with 50 mL of FBS (Hyclone)) supplemented with 10 ng/mL brain-derived neurotrophic factor (BDNF) and 10 ng/mL of neurotrophin-3 (NT-3).^62^

Organoid cultures exhibited heterogeneity in size and morphology. To mitigate this variability and ensure robust RNA-seq analysis, we initiated cultures with eight biological replicates in order to yield sufficient RNA for multiple independent ARM-seq libraries at each time point. Samples exhibiting low sequencing depth (<75k reads) were excluded from further analysis. This resulted in a larger number of biological replicates were available at earlier time points (n=8) compared to later time points (n=2).

### Immunofluorescence Microscopy

Organoids were collected and fixed in 4% Paraformaldehyde (PFA) (ThermoFisher), washed 3x with PBS, then incubated in 500 μL of 30% Sucrose for 2-3 days at 4°C until they began to float. They were then embedded in square cryomolds with Tissue-Tek O.C.T. Compound (Sakura) and placed at −80°C for storage. They were then sectioned to 18 μm using a cryostat (Leica Biosystems) directly onto glass slides. After allowing samples to come to room temperature, 3 washes of 35 minutes in 1X PBS were performed. The sections were then incubated in a blocking solution of 10% BSA for 2 hours. The sections were then incubated in primary antibodies and blocking solution at 1:1000 dilution overnight at 4°C **(Table S4)**. They were then washed 3 times for 30 minutes and incubated in secondary antibodies for 2 hours at room temperature. They were then washed 3 times for 30 minutes in PBS and sealed with ProLong™ Gold Antifade Mountant with DAPI (ThermoFisher Scientific # P36935). Microscopy was performed using a Zeiss Axio Imager and Zen Software suite, with processing of the images performed using Fiji.^63^

### RNA Isolation

Isolation of total RNA from cerebral cortical organoids and stem cells was performed using Direct-Zol RNA MiniPrep Kit (Zymo Research) with TRI Reagent (Molecular Research Center, Inc.). Since a single organoid yields far less total RNA than required for one sequencing library, approximately 3-8 organoids would be pooled in Trizol depending on organoid size for each sample replicate. The manufacturer’s recommended volume of TRI Reagent was added to each sample (∼1 mL). For RNA purification of stem cells, Trizol was directly added to cell culture plates on ice, scraped, and homogenized via pipetting. Organoids in Trizol were broken down via pipetting inside a 1mL Eppendorf tube on ice via a syringe. All total RNA was processed using a MirVana miRNA Isolation Kit (ThermoFisher Scientific), according to the manufacturer’s instructions, to select for RNA <200 nt. This was followed by an RNA Clean and Concentrate-25 (Zymo Research).

Samples were divided into plus-AlkB experimental treatment and minus-AlkB control as previously described in ARM-seq methods.^24^ AlkB sample treatment was used to effectively increase RT processivity of hyper-modified tRNA/tDRs while also removing sequencing bias favoring hypo-modified tRNA/tDRs **(Figure S1A)**. For example, the proportion of ARM-seq tRNA reads more than doubles (8.9% to 21.8%) when comparing untreated versus AlkB-treated samples. This was followed by a phenol-chloroform cleanup treatment. RNA samples were then used for library preparations.

### ARM-seq library preparation

ARM-seq libraries were constructed as described previously ^24^ utilizing the NEBNext Multiplex Small RNA Library Prep Set (New England Biolabs). Treated RNA (Minus- or Plus-AlkB treatment; 100ng total small RNA) was used as input into the library preparation, and ¼ reaction volume of all reagents were used with the NEBNext kit. PCR-amplified libraries were purified using a phenol-chloroform extraction cleanup and then size-selected (140-250 nts) on a 6% non-denaturing TBE-acrylamide gel to remove unwanted primer dimer products. Libraries were eluted from sliced gel pieces using Gel Elution Buffer (New England Biolabs) and precipitated using 0.3 M NaOAc, 80% Ethanol, and 1 μL of Linear Acrylamide (supplied in NEBNext Kit) at final concentration. Samples were left in −80°C freezer overnight to precipitate. Precipitated libraries were then pelleted, washed twice in 80% Ethanol, and resuspended in pure H2O. Libraries were then quantified using Qubit and Agilent DNA High Sensitivity kit.

### RNA sequencing and differential expression analysis

Libraries prepared for ARM-seq were sequenced using either Illumina MiSeq or Illumina NextSeq 550. 75-nt pair-ended reads were produced as FASTQ files that were analyzed using tRNA Analysis of eXpression (tRAX).^30^ Sequence adapters were trimmed, and pair-ended reads were merged using the tool trimadapters.py in tRAX. The reference database for tRAX was built with high-confidence tRNA predictions retrieved from the Genome tRNA Database^64^ and the sequences of human genome assembly GRCh38. Other gene annotations were obtained from Ensembl release 102. Biological replicates of organoids at each time point were grouped as sample replicates for tRAX inputs and different time points were marked as pairs for differential expression comparison for each sequencing type. Default options of tRAX were used, and samples with low sequence depth and outliers after DESeq2 correction were removed from analysis, ensuring duplicate at a minimum across all time points. To check for biases in organoid time points (due to unequal replicate distribution), total normalized reads **(Figure S1A,B)** were compared, showing no notable differences in abundance between time points. Additionally, we found the relative levels of tRNAs were consistent between sample groups, with no significant difference in total reads found between samples within AlkB treatment groups **(Figure S6A,B)**. AlkB treatment significantly increases tRNA read counts relative to overall RNA reads **(Figure S1B)**. ARM-seq also preferentially picks up more tRNA fragments and their read counts relative to full-length tRNAs **(Figure S1C)**. In order to standardize tRNA coverage profiles, each series of reads was aligned to conserved tRNA positions ^65^, dropping gap and extension positions. All visualizations were created using tRNAgraph (https://github.com/alba1735/tRNAgraph) and custom Python scripts (https://github.com/alba1735/organoidtRNAs).

### Northern Blot Analysis

Northern blot analysis was conducted to assess the expression levels of specific tRNAs over a 70-day time course and was modified from Damm et al. ^66^. Previously extracted day 0 and day 70 Total RNA was used. RNA samples (5-20 µg per lane) were resolved on a 15% TBE-urea denaturing polyacrylamide gel and transferred to a positively charged nylon membrane (Sigma) using a Trans-Blot® Turbo™ Transfer System (Bio-Rad). Membranes were then UV-crosslinked using an EDC (N-(3-Dimethylaminopropyl)-N’-ethylcarbodiimide)-mediated (Sigma) crosslinking protocol. Subsequently, membranes were pre-hybridized in ULTRAhyb Ultrasensitive Hybridization Buffer (Thermo Fisher) before overnight hybridization with custom 3’ biotinylated LNA probes (IDT) designed against specific tRNA species: Arg-TCT-4, Ala-AGC-8, and SeC-TCA-1 at a final concentration of 50 pmol/mL **(Table S4)**. Following hybridization, membranes were subjected to a series of stringent washes using low-stringency (2x SSC, 0.1% SDS) and high-stringency (0.1x SSC, 0.1% SDS) buffers. Detection was performed using the Chemiluminescent Nucleic Acid Detection Module (Thermo Fisher), and signal was visualized using a Chemi-Doc imager (Bio-Rad). Since specific tRNA/tDR concentration is low, a more sensitive substrate, SuperSignal™ West Femto Maximum Sensitivity Substrate (Thermo Fisher), was employed for signal detection. To ensure equal loading across lanes, the prestained DynaMarker Prestain Marker for Small RNA Plus (DiagnoCine) was used. Additionally, membranes were stained pre-transfer with SYBR™ Gold Nucleic Acid Gel Stain (Thermo Fisher) and imaged under UV light to verify the presence and integrity of the transferred RNA. Membranes were stripped using a boiling 0.1% SDS solution and reprobed as necessary following the overnight hybridization protocol described above. The Northern blots were normalized with a 5S probe for quantification.

### tRNA Classification Analysis

Sequencing output data from tRAX was used for tDR classification. DESeq2 normalized read counts were combined into an AnnData object and were arranged by coverage-associated data across human GRCh38 tRNAs.^67^ These were comprised of unique coverage (normalized read count with reads that are uniquely mapped to feature at the specific position), 3’ and 5’ read ends (with consideration to the tRNA structure), deletions (number of reads that have a gap at the specific position), and mismatched bases per position (normalized read count that does not match the reference base at specific position). Unique reads were chosen over total reads as this provided more specific coverage changes in the clusters and better tRNA uniformity therein. Thus, each cluster portrays the coverage makeup of component tDRs and gives a fragment profile in addition to other underlying sequence features. The data was further preprocessed by removing read counts less than 20, regressing out the number of reads, and scaling and centering the data. Dimensionality reduction was performed using Uniform Manifold Approximation and Projection (UMAP) and clustering using HDBSCAN.^50,51^ Classifying tRNAs into neural, neutral, and stem categories was based on the log_2_ fold-changes between day 0 and day 70 time points split into the top and bottom 20th percentiles of all tRNAs, after removing reads with low coverage and p-value >0.05. Log_2_ fold-change was normalized between −1 and 1

with each tRNA transcript and assigned a “neural stem score” based on its relative expression between the differentiated and undifferentiated time points. To determine if a cluster was neural or stem specific, the mean of the “neural stem scores’’ was taken for each cluster, and the top and bottom 25 percentiles correlated to neural (B) and stem (A) clusters, respectively. Clusters from neither group were labeled as neutral (C) and unannotated (U) results from HDBSCAN, their own group. Sequence logos were generated using Logomaker ^68^ by using the mean normalized read counts per position and converting them into information scores.^69^

### tRNA Modification Analysis

To derive tRNA modifications for analysis, we utilized data from Zhang et al. ^20^, specifically, we obtained the HEK293T cell line RNA sequencing data generated using OTTRseq, which provides full-length sequencing information necessary for modification analysis. We processed this data using the tMAP pipeline (https://github.com/Aimann/tMAP), as described in their paper. During this processing, we flagged any tRNA positions exhibiting a mismatch rate greater than 5% compared to the reference tRNA sequence. These mismatches are indicative of potential RNA modifications and, we cross-referenced the flagged tRNA positions with the Modomics database, a repository of known RNA modifications ^70^ as well as commonly cited modifications in human.^71^ Any flagged positions that matched a known modification were then labeled and projected onto the clusters (described above) identified in our primary experimental data.

## AI Statement

During the preparation of this work the authors used Grammarly and Gemini in order to restructure sentences for clarity and conciseness. After using this tool/service, the authors reviewed and edited the content as needed and take full responsibility for the content of the publication.

