## Supplemental Figures for "Identification of Specialized tRNA Expression in Early Human Brain Development"

### Supplementary Figure Legends

#### Figure S1: ARM-seq expression patterns

**(A-B)** Normalized read count distribution for tRNA reads by **(A)** AlkB treatment and **(B)** time point for ARM-seq with significance by Welch's t-test shown as ns (p <= 1), * (1e-02 < p <= 5e-02), ** (1e-03 < p <= 1e-02), *** (1e-04 < p <= 1e-03), and **** (P <= 1e-04). **(C)** Human tRNA and tDR reads-counts distributions showing relative ratio of fragments vs full-length tRNAs across organoid time course, sequencing type and AlkB treatment. **(D)** Human tRNA isotype reads-counts distributions showing relative ratio of tRNA isotypes across organoid time course, sequencing type and AlkB treatment.

#### Figure S2: Maximized fragment distributions of tDR Isotypes

**(A)** Coverage profiles for tDRs over organoid neurogenesis. D-loop, anticodon-loop, and T-loop shown in gray, with lines added at specific coverage breakpoints in each plot. The mean value was taken across all coverage profiles within a specific isotype at each position, with isodecoders not expressing a mean >= 20 reads are not displayed.

#### Figure S3: tRNA expression in neural organoids

**(A-B)** Heatmap showing the top 15 neural and top 15 stem favored tDRs across all conditions by p-value **(A)** and read counts **(B)**. P-value and normalized read counts are shown in green with a cutoff of <=0.001 and >=1000 respectively. **(C)** Coverage profiles for Arg-TCT-1 and Arg-TCT-4 ARM-seq over organoid neurogenesis. D-loop, anticodon-loop, and T-loop shown in gray, with lines added at specific coverage breakpoints in each plot. **(D)** Northern blot analysis of Arg-TCT-4, Ala-AGC-8 and SeC-TCA-1 probes at full-length mature tRNAs between day 0 and day 70 time points with 5S also provided for comparison. Lower range of the membrane was imaged with longer exposure for the tRNA probes with the upper portion of the membrane blocked, signal for the Ala-AGC-8 probe was too low to detect anything. **(E)** Coverage profiles for the tDRs that appear to downregulate with neural progenitors (D14 and D35). D-loop, anticodon-loop, and T-loop shown in gray, with lines added at specific coverage breakpoints in each plot.

#### Figure S4: tDR clustering and associated enzyme expression

**(A-B)** Bar plots of each tDR class's relative isotype **(A)** and AlkB treatment **(B)** distribution by each HDBSCAN class. **(C-D)** UMAP projections of tRNA sequencing profiles using unique coverage, 3’ and 5’ read ends (with consideration to the tRNA structure), deletions, and mismatched bases at position after HDBSCAN clustering. Projections of **(C)** tRNAscan-SE covariance and HMM as well as secondary structure (Infernal) scores and **(D)** $Log_2$ fold-change normalized expression between day 0 and day 70 time points masked by unannotated (U) class are shown. **(E)** Normalized base-mean tRNA enzyme expression from human embryonic stem cells cultured into cerebral cortical organoids on a 5 week (day 35) time course.

#### Figure S5: K-mer frequency distribution

**(A)** Sequence logo representation of the read counts for tRNAs found in neural (B) clusters dropping non-expressed tRNAs (>20 reads) and treating each sequence as if the full sequence was transcribed to prevent 5'/3' bias. **(B)** Kmer frequency plot showing the percent frequency change from background kmer frequency for neural (B), neutral (C), stem (A) and unannotated (U) clusters, after filtering out low (>15) read counts. **(C)** Sequence map of Arg-TCT tRNAs and top 15 most neural expressed tDRs.

#### Figure S6: Mean expression of small RNA reads

**(A-B)** Normalized read count distribution for tRNA reads by **(A)** AlkB treatment and **(B)** time point for ARM-seq with significance by Welch's t-test shown as ns (p <= 1), * (1e-02 < p <= 5e-02), ** (1e-03 < p <= 1e-02), *** (1e-04 < p <= 1e-03), and **** (P <= 1e-04).

### Supplementary Table Legends

#### Table S1: Differential Expression of Small RNAs

List of small RNAs detected in the ARM-seq data, with associated read counts, $log_2$ fold-changes and adjusted p-values for developmental time points.

#### Table S2: ARM-seq Differential Expression of tRNAs

List of tRNAs and tDRs detected in the ARM-seq data, with associated read counts, $log_2$ fold-changes and adjusted p-values for developmental time points of AlkB+ and AlkB- treatments.

#### Table S3: tRNAs Classes and tRNAs therein

List of tRNA clusters that were manually annotated from UMAP/HDBSCAN clustering.
Associated metadata, isotype, amino group, organoid developmental time point, AlkB treatment, number of read counts, reference tRNA gene.

**Table S4: DNA Oligonucleotides and antibodies**

List of primary and secondary antibodies concentrations used for IF-staining and LNA probes used for Northern blot analysis.

**Supplementary Figure S1**

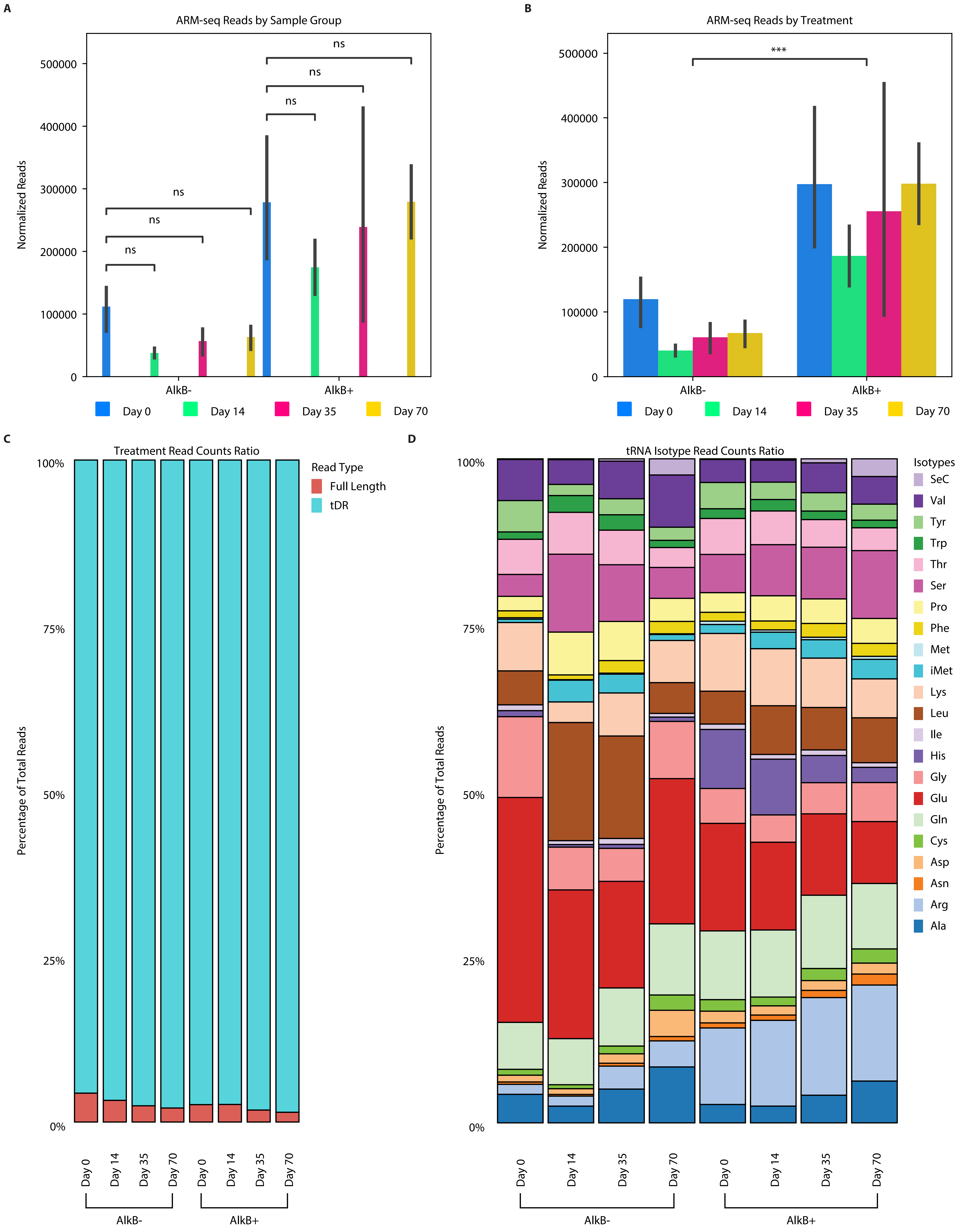

**Supplementary Figure S2**

**A**

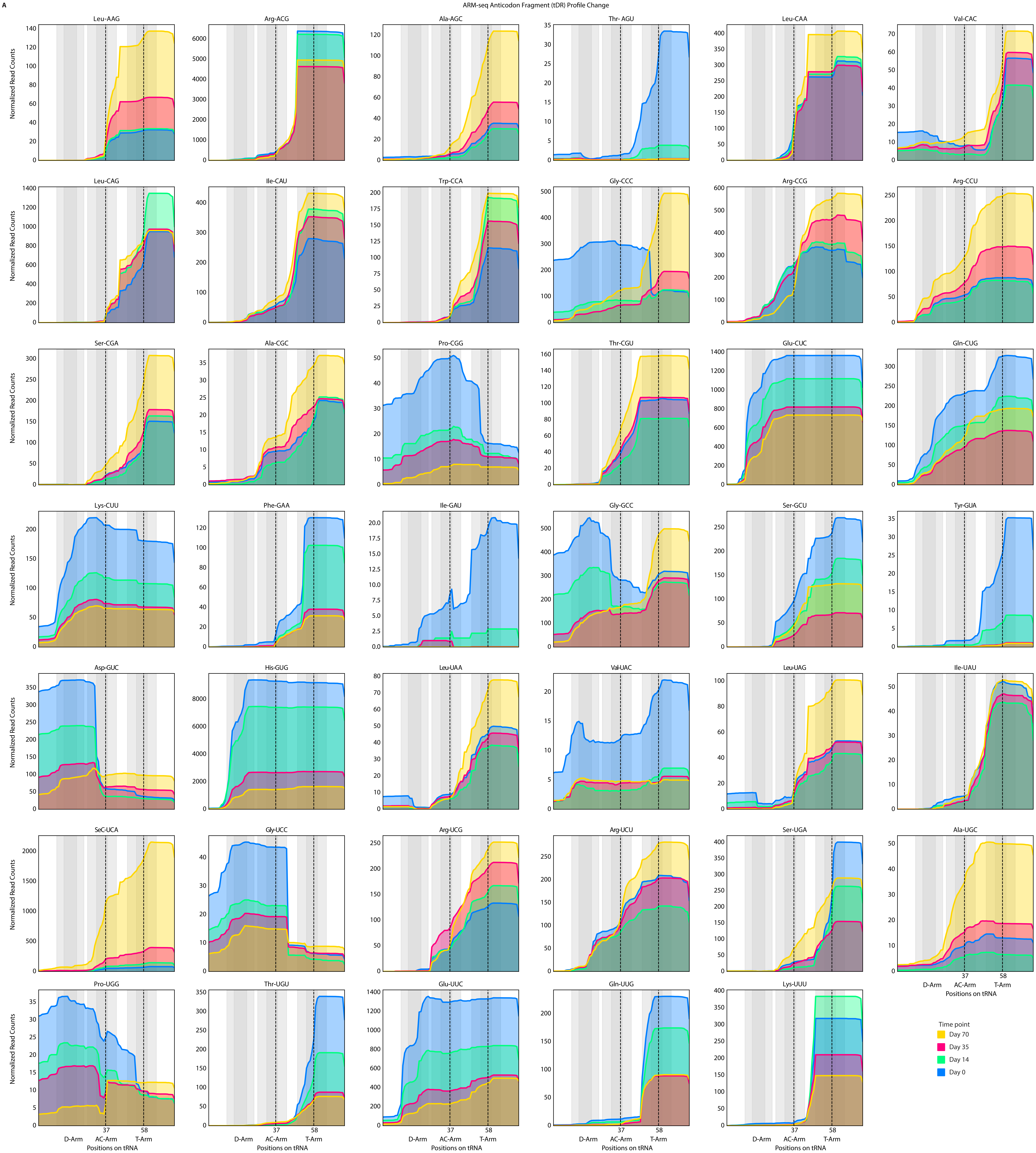

**Supplementary Figure S3**

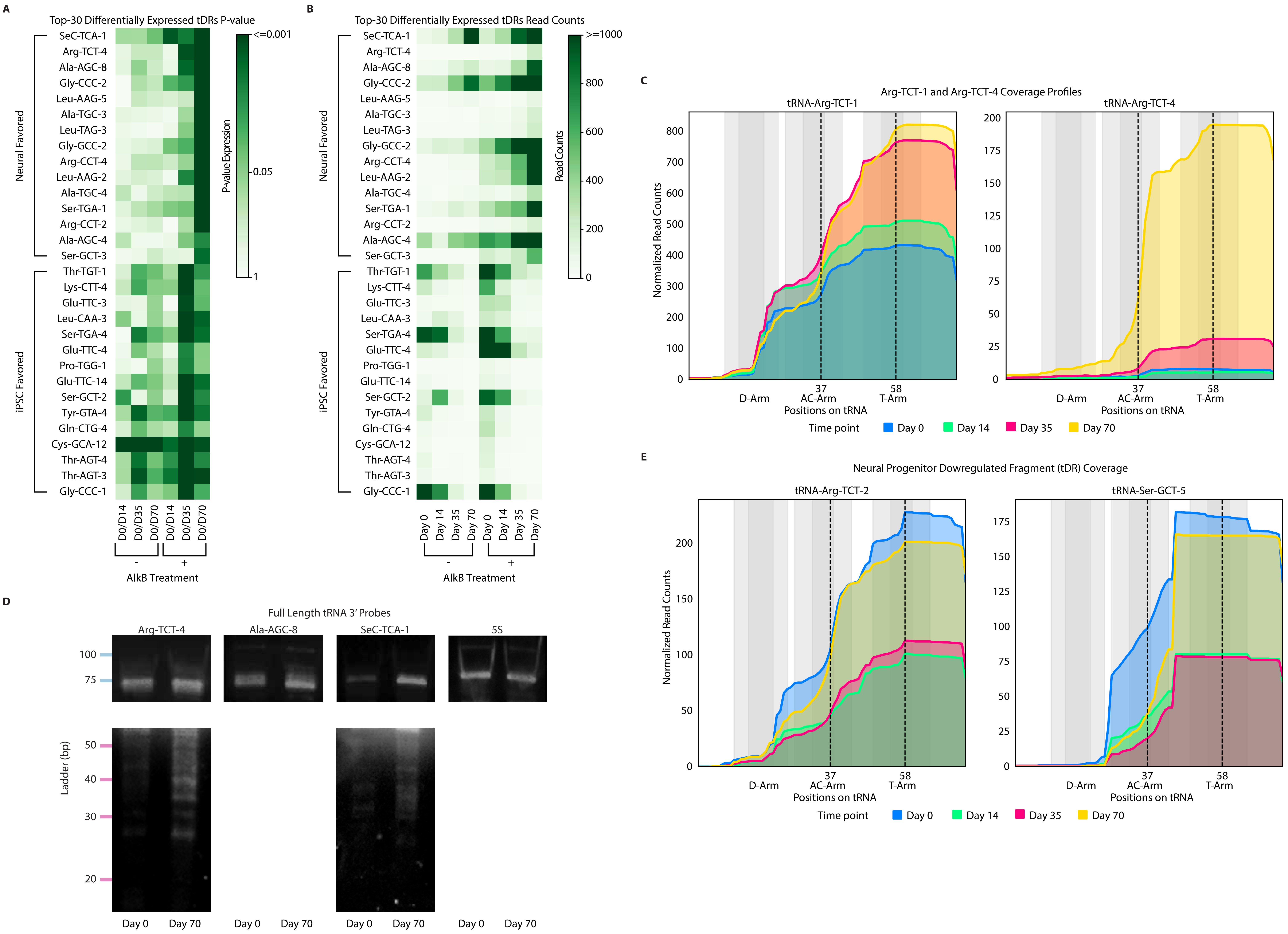

**Supplementary Figure S4**

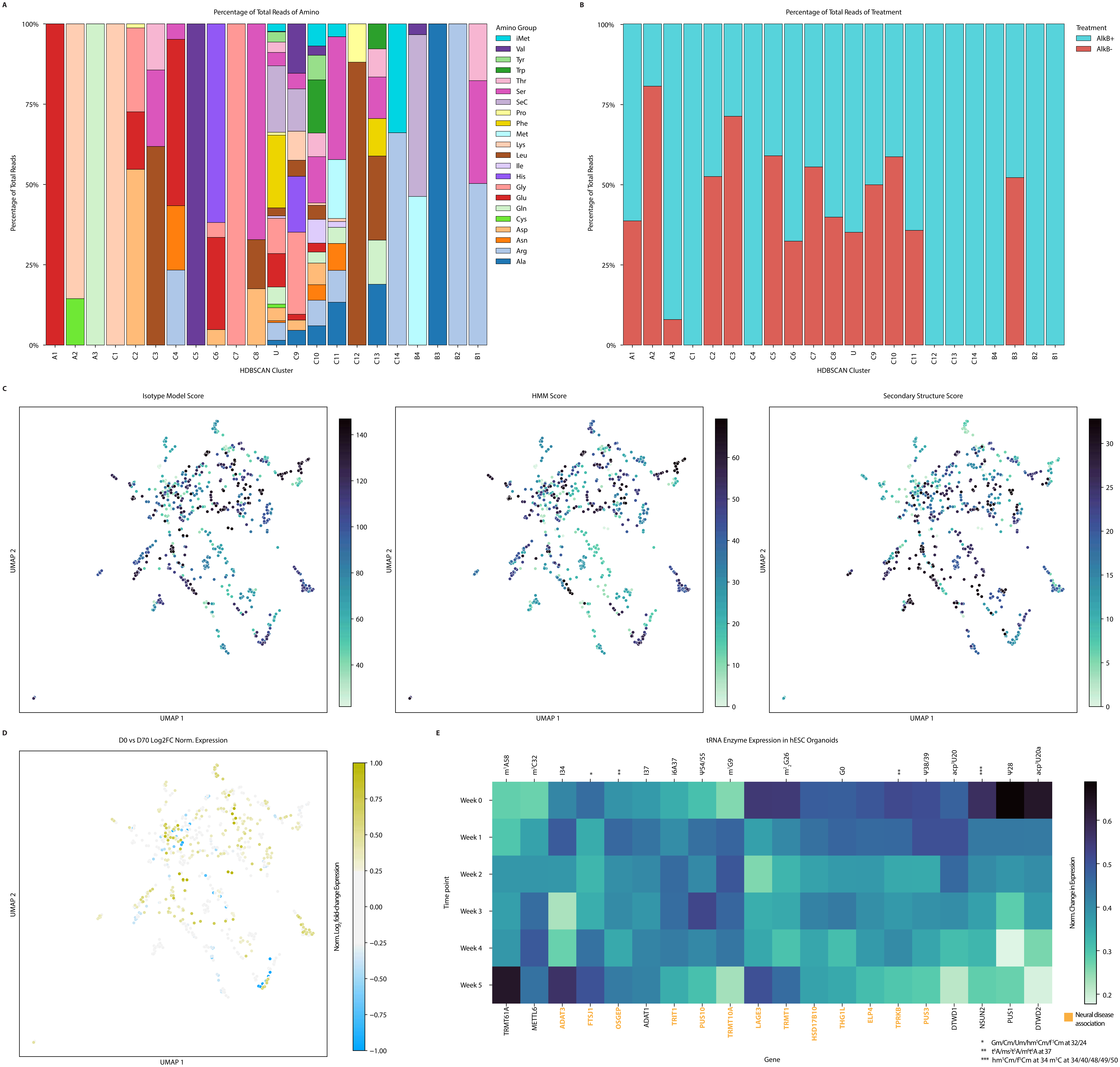

**Supplementary Figure S5**

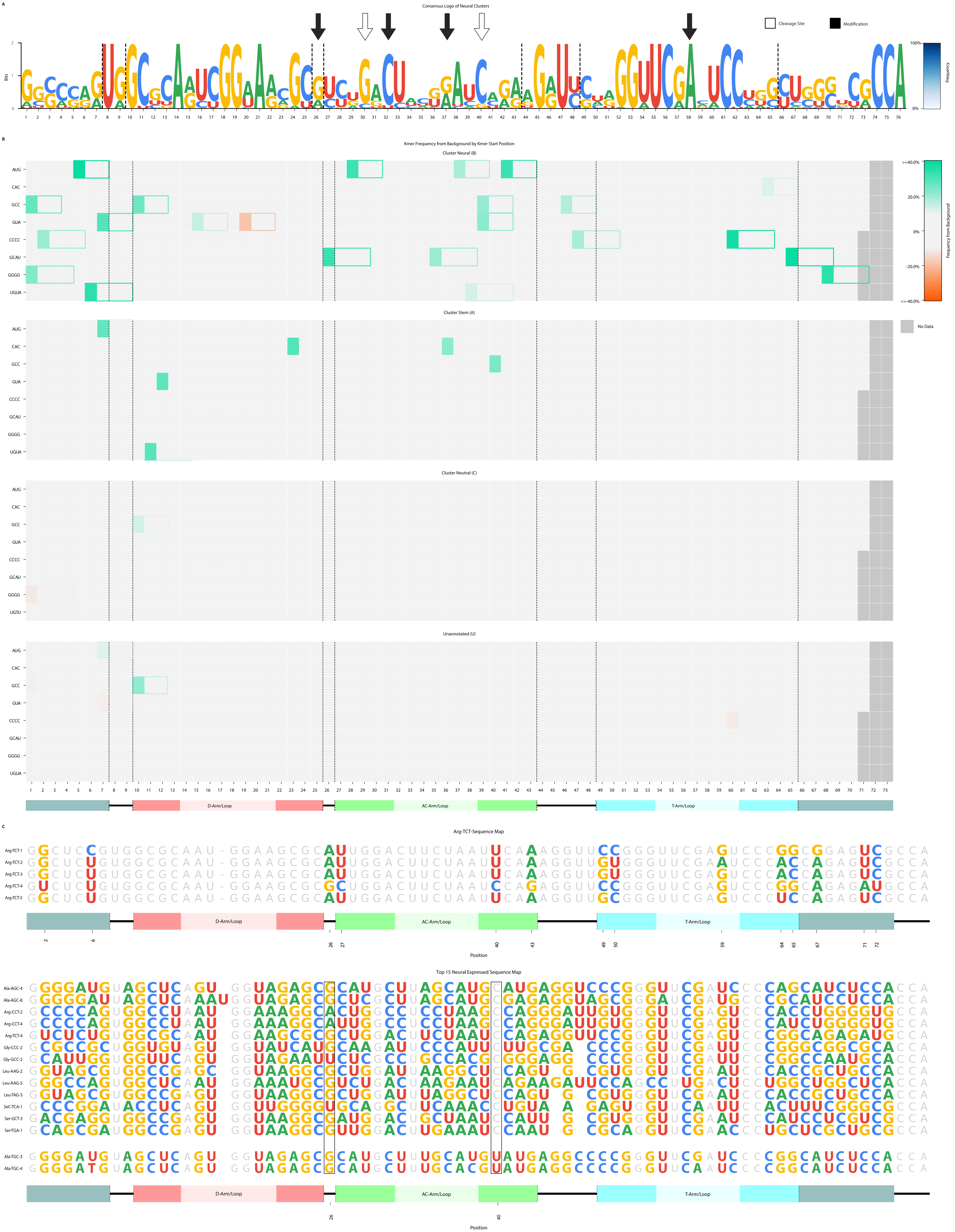

Supplementary Figure S6

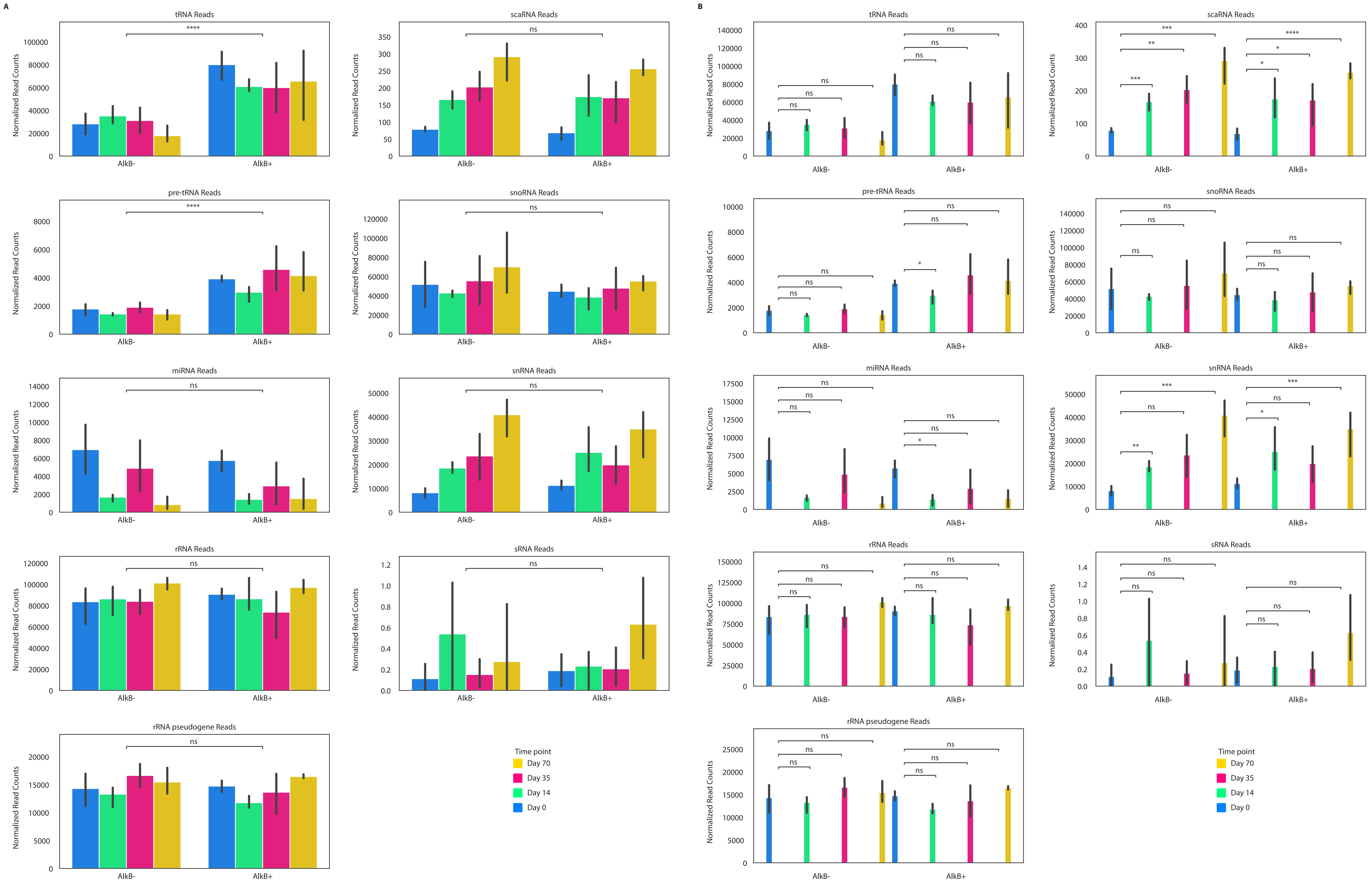
